# Evolutionary Perspectives on Anxiety: Telencephalic Circuitry and the Anxiogenic Role of TrkB Signaling in Tuberous Sclerosis Complex

**DOI:** 10.1101/2025.06.23.661056

**Authors:** Olga Doszyn, Justyna Zmorzynska

## Abstract

Tuberous Sclerosis Complex (TSC) is a genetic disease which manifests as a range of neurological symptoms, including benign brain tumors, epilepsy, and TSC-associated neuropsychiatric disorders (TANDs). Among the latter, according to recent reports, anxiety and mood disorders affect over 50% of patients. We have previously demonstrated anxiety-like behavioral symptoms in the zebrafish model of TSC, which were rescued by treatment with the TrkB antagonist ANA-12. Here, we aimed to investigate the mechanism of how ANA-12 regulates behavior by analyzing brain activity in the telencephalon of TSC zebrafish larvae, and we identified the affected regions as corresponding to the known mammalian circuitry involved in anxiety processing. Due to differences in development, the identification of telencephalic territories that are homologous between zebrafish and mammals remains challenging, particularly at early, dynamic stages of development. However, we were able to identify populations of neurons in the zebrafish habenula and ventral subpallium whose involvement in anxiety parallels that of mammals. Those regions were dysregulated in the TSC mutant. This dysregulation correlated with aberrant anxiety behavior and was rescued by treatment with ANA-12. Our results suggest that hyperactivation of TrkB in those regions is a major contributor to anxiety-like behavior as seen in TSC fish, and that those mechanisms could be evolutionarily conserved between zebrafish and mammals.

## Introduction

Tuberous sclerosis complex (TSC) is a genetic disease caused by inactivating mutations in either the *TSC1* or *TSC2* gene. The products encoded by these genes, known as Hamartin and Tuberin respectively, together with the protein TBC1D7, form the TSC complex, acting as a negative regulator of mammalian/mechanistic target of rapamycin complex 1 (mTORC1) (Crino et al., 2006). mTORC1 is a hub for many signaling pathways, and in neuronal development, it regulates the processes of axon genesis and guidance, dendritogenesis, synaptic plasticity, learning and memory (Switon et al., 2017). In the brain, TSC presents with benign tumors, lesions and cortical dysplasia, epilepsy, and TSC-associated neuropsychiatric disorders (TANDs) (Crino et al., 2006; Salussolia et al., 2019). This term is used to describe a wide plethora of symptoms, ranging from intellectual disability and cognitive deficits, autism spectrum disorder, attention deficit– hyperactivity disorder (ADHD), to anxiety and mood disorders (de Vries et al., 2015). According to the recent statistics presented by the TANDem project, which aimed to establish an internationally standardized methodology for diagnosis and treatment of TANDs, anxiety and mood disorders affect up to 56% of TSC patients. At the same time, anxiety is believed to be underdiagnosed, especially when co-occurring with intellectual disabilities, which may make it difficult for the patient to effectively communicate their experiences. The TANDem project also concludes that more research is still needed to understand the underlying mechanisms by which anxiety arises in TSC patients, and find targeted treatments (de Vries et al., 2023).

In the previous study conducted by our group, we have found elevated cortisol levels and anxiety-like behavior exhibited by *tsc2^vu242/vu242^* (tsc2-deficient) zebrafish larvae. The increased high-range velocity in *tsc2^vu242/vu242^* fish, indicative of anxiety, was rescued by treatment with ANA-12, a selective TrkB inhibitor. ANA-12 also rescued the seizure-associated decreased activity of the *tsc2^vu242/vu242^* mutant, as well as the observed thinning of the anterior commissure, suggesting the role of TrkB signaling in brain connectivity and epileptogenesis, either of which could contribute to anxiety (Kedra et al., 2020). However, the question of how ANA-12 might affect the activity of pertinent brain regions in order to produce its effect on behavior remained unanswered.

In this study, we conducted an in-depth investigation into the anxiolytic effects of ANA-12 by performing additional behavioral assays in zebrafish. Furthermore, we examined neuronal activation patterns in the frontbrain of tsc2^vu242/+^ zebrafish larvae using whole-brain imaging techniques to elucidate the neuroanatomical correlates of ANA-12-induced behavioral changes and we have found that ANA-12 regulates activity in regions linked to behavioral regulation and processing of stress and fear related information, such as the habenulae, striatum and septum. We have also putatively identified parts of the amygdaloid complex in the zebrafish brain at early stages of development.

## Methods

### Zebrafish breeding and genotyping

All experiments were performed using the *tsc2^vu242/+^* zebrafish line (Kim et al., 2011). Adult and larval zebrafish were maintained according to international standards. In each experiment, offspring of at least three parental pairs were used. The larvae were genotyped in order to distinguish wild-type fish from those hetero- or homozygous for the mutation in the *tsc2* gene. Behavioral experiments were performed blindly for the genotype, with larvae collected for genotyping afterwards; for the purpose of immunofluorescence staining, Western blot and ELISA assays, heads were collected for the experiment, and tails were used for genotyping. All genotyping was performed using the HRM method.

### Drug treatments

All stock solutions of drugs used in this study were prepared in E3 medium (5 mM NaCl, 0.17 mM KCl, 0.33 mM CaCl_2_, 0.33 mM MgSO_4_) or dimethyl sulfoxide (DMSO; Sigma-Aldrich (Merck)). Working solutions were prepared in E3 medium. Drugs were administered into the bathing medium, containing up to 50 dechorionated larvae. Treatments included: 200 nM rapamycin from 2 days post-fertilization (dpf), and 50 nM ANA-12 or 60 μM vigabatrin (all from Sigma-Aldrich (Merck)) 24 hours before the behavioral test.

### Behavioral tests

Behavioral tests were performed using zebrafish larvae at 5 dpf, following a 15-minute period of habituation to the behavioral testing room. Zebrafish activity was recorded using the ZebraBox system, with experimental parameters set using the dedicated ZebraLab software (both from ViewPoint Behavior Technology). Obtained raw data was processed and analyzed using RStudio software (cran.r-project.org; rstudio.com).

The open field test was performed as previously described (Kedra et al., 2020). Single larvae were placed in each well of a 6-well plate, filled with E3 medium. The plate was uniformly illuminated with bottom light set to 80% intensity. Activity was tracked for 8 min, with a 5 s time bin. Surround and central areas of each well were defined in ZebraLab software, and the cumulative activity in each area (normalized to the total time of movement) was calculated for each fish. Lack of movement of the *tsc2^vu242/vu242^* mutants was mapped to non-motor seizures before (Kedra et al., 2020), therefore not moving fish were excluded from the analysis.

For the sudden light changes test (Kedra et al., 2020), single larvae were placed in each well of a 24-well plate, filled with E3 medium. Activity was tracked for 30 min overall, with a 5 s time bin. The experiment consisted of three phases: dark-light-dark, of 10 min each. During the light phase, the plate was illuminated with bottom light set to 60% intensity.

### Whole-mount immunofluorescence

At 5 dpf, heads of zebrafish larvae were collected into 4% PFA solution for overnight fixation at 4°C. For staining with antibodies against phosphorylated proteins, 2% (v/v) of 1 M NaF was added to the fixative solution as a phosphatase inhibitor. Fixed samples were then washed with 1x PBS, and incubated with 3% KOH, 1% H_2_O_2_ bleaching solution for approx. 30-45 min at room temp. to remove pigmentation. Samples were then stained with primary antibodies (anti-pRps6 (Ser235/236; Cell Signaling Technology), 1:200; anti-TrkB (Proteintech), 1:400; anti-ERK (Cell Signaling Technology), 1:400; anti-pERK (Cell Signaling Technology), 1:400; anti-VGlut1 (Synaptic Systems), 1:400; GAD65/67 (Abcam), 1:400; anti-calretinin (Swant), 1:200; anti-NPY (Abcam), 1:200; anti-parvalbumin (Genetex), 1:250) and secondary antibodies (donkey anti-mouse Alexa Fluor 568, 1:1000, or donkey anti-rabbit, Alexa Fluor 488, 1:1000, both from Thermo Fisher Scientific) according to a previously established protocol (Doszyn et al., 2025).

### Protein extraction and Western blot

25-30 heads of each genotype were pooled into Ringer’s solution supplemented with protease and phosphatase inhibitors (1 mM aprotinin, 1 mM leupeptin, 0.25 mM benzamidine hydrochloride, 0.25 mM Pefabloc® SC, 0.5 mM sodium orthovanadate, 1 mM sodium β-glycerophosphate, 0.25 mM tetrasodium pyrophosphate, 2.5 mM sodium fluoride), 1% Triton™ X-100 (Sigma-Aldrich (Merck)), 0.1% SDS, and 0.5 mM EDTA, sonicated, and placed on ice for 30 min. The samples were then centrifuged for 5 min at 4 ℃ at 9000 rpm, and concentrated Laemmli buffer was added to each sample to the final concentration of 1x. The samples were then denatured by incubation in 95 ℃ for 5 min using Thermomixer C (Eppendorf®), aliquoted and stored at −20 ℃.

SDS-PAGE electrophoresis was performed using the vertical Mini-PROTEAN system (Bio-Rad), in Tris-Glycine-SDS buffer. For the immunodetection of target proteins, the samples were transferred from the polyacrylamide gel onto a PVDF membrane (Sigma-Aldrich (Merck)) activated by submerging in 100% methanol. Transfer was performed using the vertical Mini Trans-Blot® Cell system (Bio-Rad), in Towbin buffer with SDS, at 4 ℃. The membrane was then washed with deionized water, and incubated in 5% milk in TBS buffer against non-specific binding for 1 h at room temp. After blocking, the membrane was washed with TBST, and incubated with primary antibodies: anti-Rps6 (Santa Cruz Biotechnology), 1:200; anti-pRps6 (Ser235/236 Cell Signaling Technology), 1:1000) in 5% BSA in TBST overnight at 4 ℃. On the following day, the membrane was washed thrice with TBST, and incubated with secondary antibodies (IRDye® 680RD Donkey anti-Rabbit IgG, 1:10 000, or IRDye® 800CW Donkey anti-Mouse IgG, 1:10 000, both from LI-COR Bioscience) in 5% BSA in TBST for 1 h at room temp. Afterwards, the membrane was again washed thrice with TBST, then imaged using the Odyssey DLx (LI-COR Bioscience), and analyzed using Image Studio Lite v.5.5 software (LI-COR Bioscience).

### ELISA assay

The ELISA assays for P-TrkB and TrkB were performed on samples containing 20 heads each, according to manufacturers’ instructions. The detection antibody in both assays was the biotin-conjugated anti-TrkB antibody (500 ng/ml working concentration; catalog no. BAF397, R&D) and streptavidin-horseradish peroxidase (1:200; R&D) to detect TrkB. The capture antibodies were anti-P-TrkB (catalog no. DYC688, R&D) and anti-TrkB (4 ng/l working concentration; catalog no. 13129-1-AP, ProteinTech).

### Imaging and image analysis

All images were acquired using a Zeiss Lightsheet Z.1 microscope (40x water immersion objective, NA = 1.3) at 1024 × 1024 pixel-resolution, in a z-stack mode with an interval of 0,5 μm between z-slices, resulting in multi-layered images of brain tissue.

The images were then analyzed using Fiji software (Schindelin et al., 2012). For calretinin, the number of calretinin-positive cells in each region of interest (ROI) was counted manually. For phosphorylated ribosomal protein s6 (pRps6), 5 signal-positive cells per fish were randomly selected, mean signal intensity of each cell were measured using the Measure tool, and the mean intensity per fish was later calculated in Rstudio. For TrkB and parvalbumin, ROIs were selected manually, Z-slices were superimposed using the Sum slices tool, and the intensity or integrated density of the fluorescent signal were measured using the Measure tool. Integrated density was chosen as a proxy for the number of parvalbumin-positive cells due to the low optic resolution in the medial subpallium, making it difficult to distinguish single cells.

For, neuronal activity measurements using pERK-ERK method, the brains were firstly registered to the reference brain using Parallel Fiji CMTK Registration Plugin by Sandor Kovacs (github.com/sandorbx/Parallel-Fiji-CMTK-Registration). After accuracy analysis, the automated analysis of neuronal activity was performed similarly to previously described protocol (Randlett et al., 2015), except the commands were done in Fiji. These included preprocessing, division of pERK and ERK channels, gaussian blurring, and averaging. At least 20 images were averaged per genotype per treatment from two independent experiments. The statistics were calculated using FDR method versus randomized dataset. Only significant pixels are represented on the neuronal activity maps (images).

### Statistical analysis

All data were analyzed with Rstudio software ((cran.r-project.org; rstudio.com). Sample sizes could not be predetermined due to the random distribution of genotypes; however, this provided randomization and blinding for sample collection. Equality of variance and normality of residuals were determined with Levene’s test and Shapiro-Wilk test, respectively. If data were normally distributed, they were analyzed by two-way ANOVA with post-hoc TukeyHSD test; otherwise, the Kruskal-Wallis test was used with post-hoc Wilcoxon test to correct for multiple comparisons. Data are presented as medians using boxplots, where each dot represents datapoint from one fish. Data points outside of the boxplot whiskers represent outliers. Adjusted p-values below 0.05 were considered statistically significant, and are shown on figures with * for p < 0.05, ** for p < 0.01, *** for p < 0.001, and **** for p < 0.0001.

## Results

### ANA-12 rescues anxiety-like behavior in tsc2^vu242/vu242^ fish

We have previously shown that the *tsc2^vu242/vu242^* fish exhibit hypervelocity, thigmotaxis in the open field test, and anxiety-like behavior in response to a sudden changes in light conditions. Additionally, hypervelocity was rescued by treatment with ANA-12 (Kedra et al., 2020). Here, we have performed the open field and sudden light changes tests using *tsc2^vu242^* fish pretreated with ANA-12 versus untreated, and have confirmed a rescue of anxiety-like behaviors in those tests by ANA-12. Relative time spent next to the walls of the testing chamber was increased in *tsc2^vu242/vu242^* fish in comparison to wild-type *tsc2^+/+^* and heterozygous *tsc2^vu242/+^* siblings representing anxiety-like behavior. This was reduced in *tsc2^vu242/vu242^* homozygotes following treatment with ANA-12 (Fig.1 A-C). However, a reduction in thigmotaxis was also observed following pretreatment with the mTorC1 inhibitor rapamycin, and the antiepileptic drug vigabatrin (Fig.1 B-C). Mutant *tsc2^vu242/vu242^* fish also showed hyperactivity during the dark phases of the sudden light changes test, which is indicative of anxiety, although freezing during the light phase was not increased in relation to wild-type *tsc2^+/+^* siblings, whose activity levels also dropped drastically when the light was switched on (Fig.1 E). The hyperactivity of *tsc2^vu242/vu242^* during the dark phase was markedly reduced by treatment with ANA-12 or rapamycin (Fig.1 F-H).

**Figure 1.**
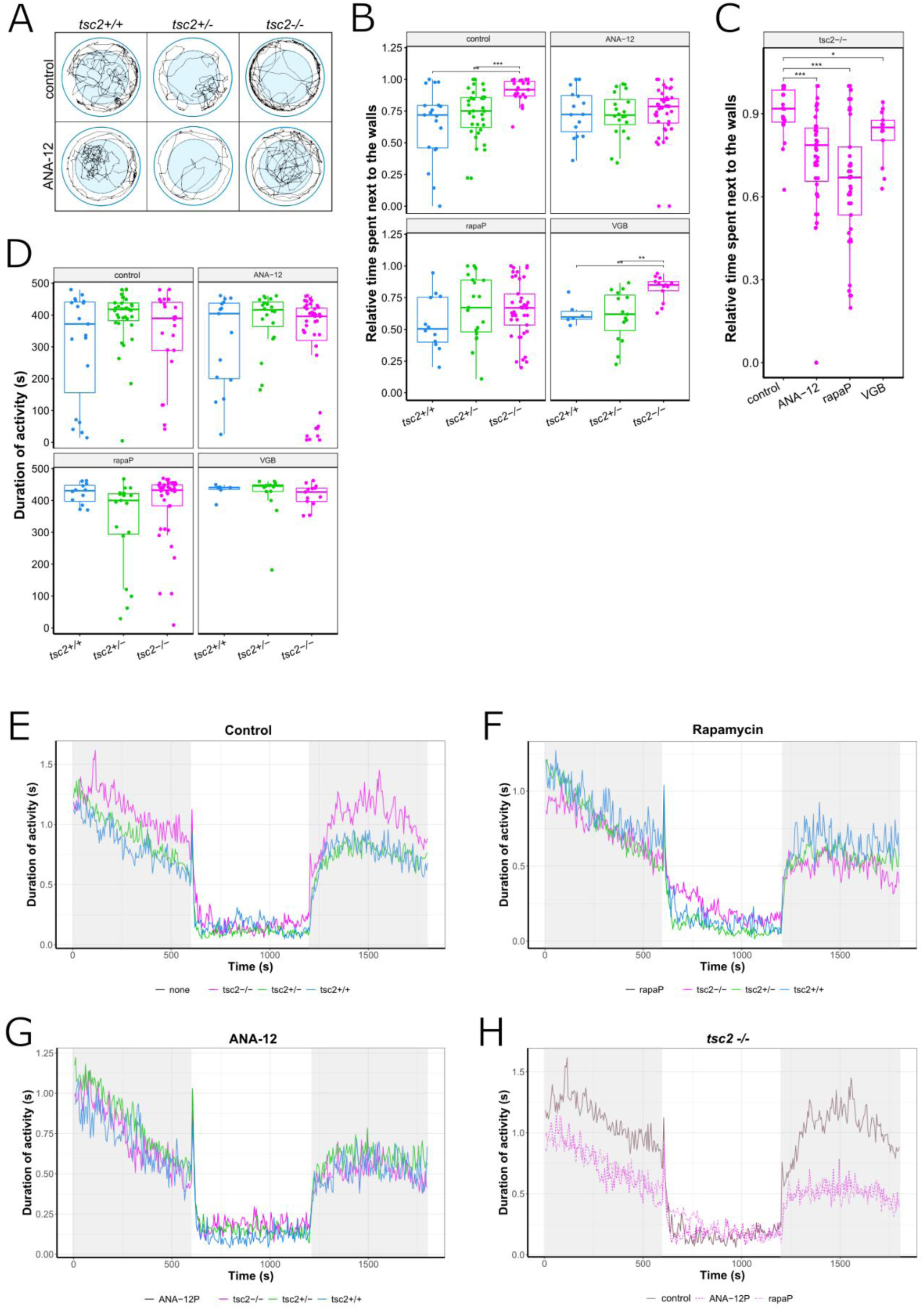
*Tsc2^vu242/vu242^* fish show anxiety-like behavior that is ameliorated by pretreatment with ANA-12. RapaP – rapamycin pretreatment starting at 2 dpf in order to prevent the development of symptoms; VGB – vigabatrin. (A) Representative tracks from the open field test for each *tsc2^vu242^* genotype in the control group vs. treated with ANA-12. Summary tracks from 8 min recordings show thigmotaxis behavior in *tsc2^vu242/vu242^* fish that is rescued by ANA-12 treatment. (B) Relative time spent next to the walls of the dish [*p* = 0.002 for control *tsc2^+/+^* vs. control *tsc2^vu242/vu242^*, *p* = 0.000159 for control *tsc2^vu242/+^* vs. control *tsc2^vu242/vu242^*, *p* = 0.002 for VGB *tsc2^+/+^* vs. VGB *tsc2^vu242/vu242^*, *p* = 0.002 for VGB *tsc2^vu242/+^* vs. VGB *tsc2^vu242/vu242^*]. (C) Relative time next to the walls compared between *tsc2^vu242/vu242^* fish only [*p* = 0.000227 for control vs. ANA-12, *p* = 0.000122 for control vs. rapamycin]. (D) Duration of overall activity across all genotype and treatment groups. No significant difference between median activity of each group indicates that the relative activity differences are not caused by changes in overall motor activity. (E-G) Mean activity over time during suddenly changing light conditions (dark-light-dark) for control, rapamycin and ANA-12 treated groups respectively. (H) Mean activity over time during suddenly changing light conditions compared between *tsc2^vu242/vu242^* fish only.

### ANA-12 treatment affects TrkB activation

First, we checked how treatment with ANA-12 affected the levels and activity of TrkB in the whole fish heads by ELISA assay to confirm ANA-12 actions. TrkB undergoes self-dimerization and autophosphorylation following the binding of BDNF (Huang & Reichardt, 2003). As we have demonstrated before (Kedra et al., 2020), *tsc2^vu242/vu242^* fish show increased levels of phosphorylated TrkB (pTrkB) in comparison to wild-type siblings. This effect was rescued by treatment with the direct TrkB antagonist ANA-12 (Fig.2 A). The immunofluorescence stainings of the intact zebrafish brains demonstrated that there was no significant difference between genotypes in either the amount or distribution of the TrkB protein in the pallium of *tsc2^vu242^* fish, and that those parameters were also not affected by ANA-12 treatment (Fig.2 B-C).

**Figure 2.**
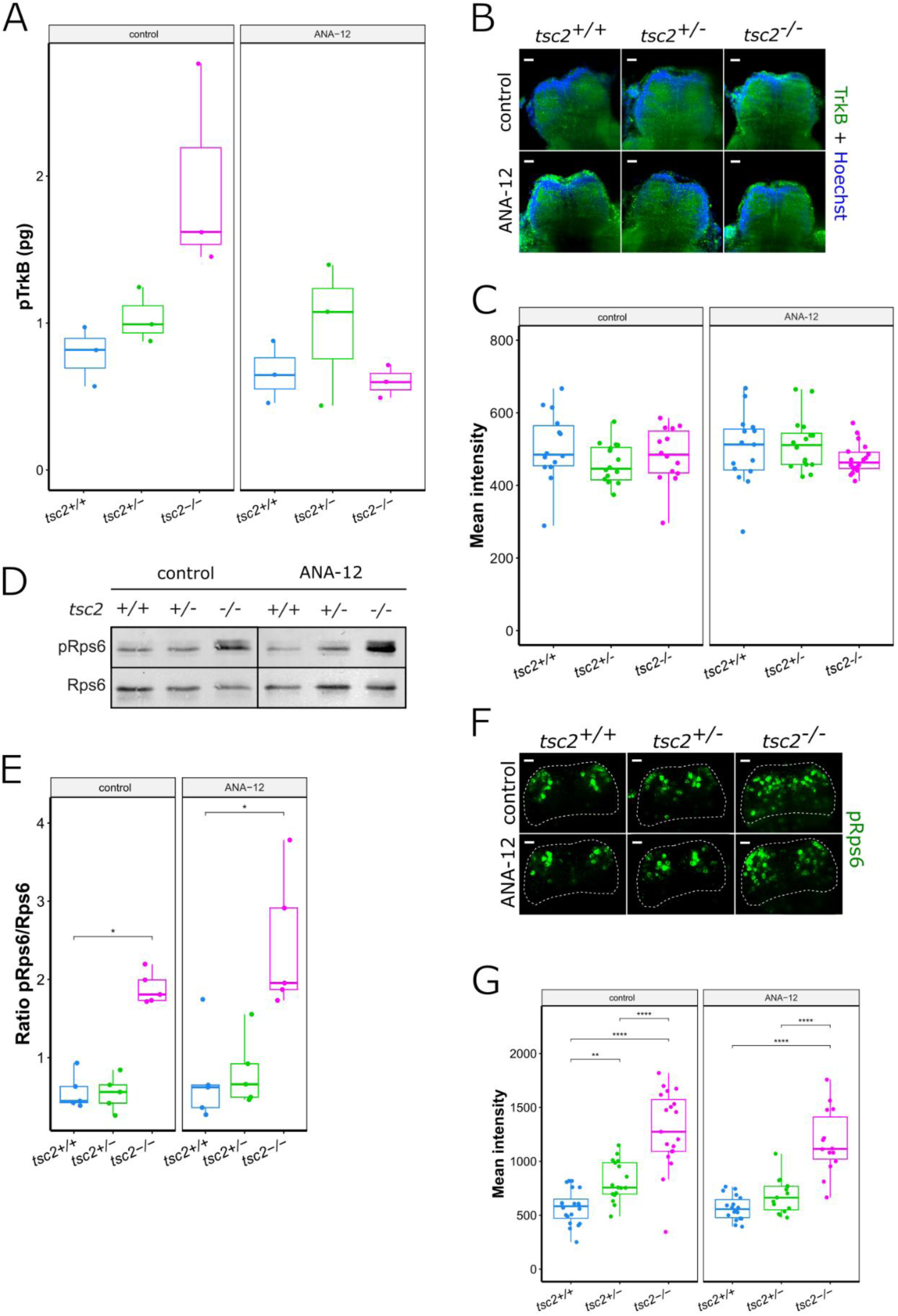
ANA-12 affects TrkB activation but not TrkB levels or mTorC1 activation. (A) pTrkB levels in *tsc2^vu242^* fish ± ANA-12 as measured by ELISA assay. (B) Representative images of the *tsc2^vu242^* zebrafish pallium stained with anti-TrkB antibody and Hoechst nuclear stain. Scale bar = 20 µm. (C) Mean intensity of TrkB signal in the pallium across all genotypes ± ANA-12. Each dot represents one fish. (D) Representative Western blot results showing the amount of pRps6 and Rps6 protein in whole brains of *tsc2^vu242^* fish ± ANA-12. (E) Ratio of pRps6 to Rps6 protein in *tsc2^vu242^* fish ± ANA-12 as measured by Western blot [*p* = 0.016 for control *tsc2^+/+^* vs. control *tsc2^vu242/vu242^*, *p* = 0.032 for ANA-12 *tsc2^+/+^* vs. ANA-12 *tsc2^vu242/vu242^*]. Each dot represents one sample pooled from at least 25 fish. (F) Representative images of the *tsc2^vu242^* zebrafish pallium stained with anti-pRps6 antibody. Scale bar = 20 µm. (G) Mean intensity of pRps6 signal in the pallium across all genotypes ± ANA-12 [*p* = 0.00399 for control *tsc2^+/+^* vs. control *tsc2^vu242/+^*, *p =* 3.636e-07 for control *tsc2^+/+^* vs. control *tsc2^vu242/vu242^*, *p =* 8.52e-05 for control *tsc2^vu242/+^* vs. control *tsc2^vu242/vu242^*, *p =* 3.054e-07 for ANA-12 *tsc2^+/+^* vs. control *tsc2^vu242/vu242^*, *p =* 4.206e-05 for ANA-12 *tsc2^vu242/+^* vs. control *tsc2^vu242/vu242^*]. Each dot represents an average intensity of five cells from one fish.

### ANA-12 treatment does not affect mTORC1 activity

The stimulation of TrkB by BDNF triggers the activation of multiple downstream pathways, including the phospholipaseCγ (PLCγ), mitogen-activated protein kinase/extracellular signal-regulated protein kinase (MAPK/ERK), and phosphatidylinositide 3-kinase (PI3K)-Akt-mTOR pathway (Huang & Reichardt, 2003; Takei et al., 2004). Although there is no functional TSC complex in the *tsc2^vu242/vu242^* fish, we checked whether ANA-12 could act through an unknown mechanism that impacts mTORC1 activity. We performed a Western blot assay to check the ratio of phosphorylated ribosomal protein s6 (P-Rps6) to non-phosphorylated form of this protein (Rps6) as it is one of the downstream targets of mTORC1. In the untreated *tsc2^vu242/vu242^* fish, the ratio of P-Rps6 to Rps6 was increased in comparison to wild-type *tsc2^+/+^* and heterozygous *tsc2^vu242/+^* siblings, owing to mTORC1 hyperactivity; and this ratio was preserved in the group treated with ANA-12 (Fig.2 D-E). Similarly, whole-mount immunofluorescence staining against P-Rps6 showed no effect of ANA-12 on neither the levels of P-Rps6 nor the distribution of P-Rps6 positive cells in the pallium of *tsc2^vu242^* fish (Fig.2 F-G). Therefore, even though pretreatment with rapamycin resulted in a similar rescue of behavioral symptoms as ANA-12, suggesting that both TrkB and mTORC1 signaling play a role in regulating anxiety, we have confirmed that the effect of ANA-12 on zebrafish behavior is not exerted directly through the mTORC1 pathway.

### ANA-12 affects brain activity

The habenula (Hb), a bilateral structure located within the epithalamus, is split into two main subdivisions – medial and lateral in mammals, corresponding to dorsal and ventral in zebrafish respectively (Amo et al., 2010). In both zebrafish and mammals, the habenula is known to play a role in a multitude of processes and behaviors, including fear and anxiety, aversion and reward, sleep and circadian rhythm, reproductive and aggressive behaviors, and processing of sensory stimuli (Fore et al., 2018).

Previously, we have found hyperactive cells in the left dorsal habenula (LDHb) of larval zebrafish, which were also marked by hyperactivation of mTORC1. This hyperactivation was linked to aberrant processing of light stimuli. We have shown that pretreatment with the mTORC1 inhibitor rapamycin rescued both the behavioral symptoms and the dysregulation of neuronal activity (Doszyn et al., 2024). Here, we have imaged the whole left and right habenula following whole-mount staining with antibodies against extracellular signal-regulated kinase (ERK) and phosphorylated (activated) ERK. Since ERK is phosphorylated in response to the calcium influx following neuron depolarization, the ratio of pERK/ERK can be used as a readout of neuronal activity, and has been validated as such in zebrafish (Randlett et al., 2015). We have calculated the ratio of pERK/ERK in untreated *tsc2^vu242^* fish and siblings treated with ANA-12, and created maps of neuronal activity in each group. Next, we have compared the composite difference maps of activity between *tsc2^vu242/vu242^* mutant *vs*. wild-type and between *tsc2^vu242/vu242^* treated with ANA-12 *vs*. untreated *tsc2^vu242/vu242^* in order to check what brain regions are hypo- or hyperactive in the mutant fish, and whether ANA-12 affect these differences. As seen before in calcium imaging experiments (Doszyn et al., 2024), we have observed significant hyperactivity in the dorsal part of the left habenula. Interestingly, ANA-12 lowered the activity in this region, even though it did not affect light preference behavior (Doszyn et al., 2024), indicating that the effect of ANA-12 might be restricted to anxiety-like behaviors, even when the affected brain regions have other functions as well. Furthermore, ANA-12 treatment did not result in uniform dampening of neuronal activity, since some regions were unaffected by the treatment, and those that were hypoactive in the *tsc2^vu242/vu242^* mutant, such as in the right dorsal habenula, were hyperactivated by ANA-12 (Fig.3 A). A similar effect was observed around the habenula commissure, which appeared separated into two distinct hyper- and hypoactive tracts, whose levels of activity were reversed after ANA-12 treatment (Fig.3 B). In the ventral part of both left and right habenula, too, cells that were hypoactive in untreated *tsc2^vu242/vu242^* fish were more active after ANA-12, and vice versa (Fig.3 C). Therefore, we demonstrated that ANA-12 could rescue both hypo- and hyperactivity in dysregulated brain regions.

**Figure 3.**
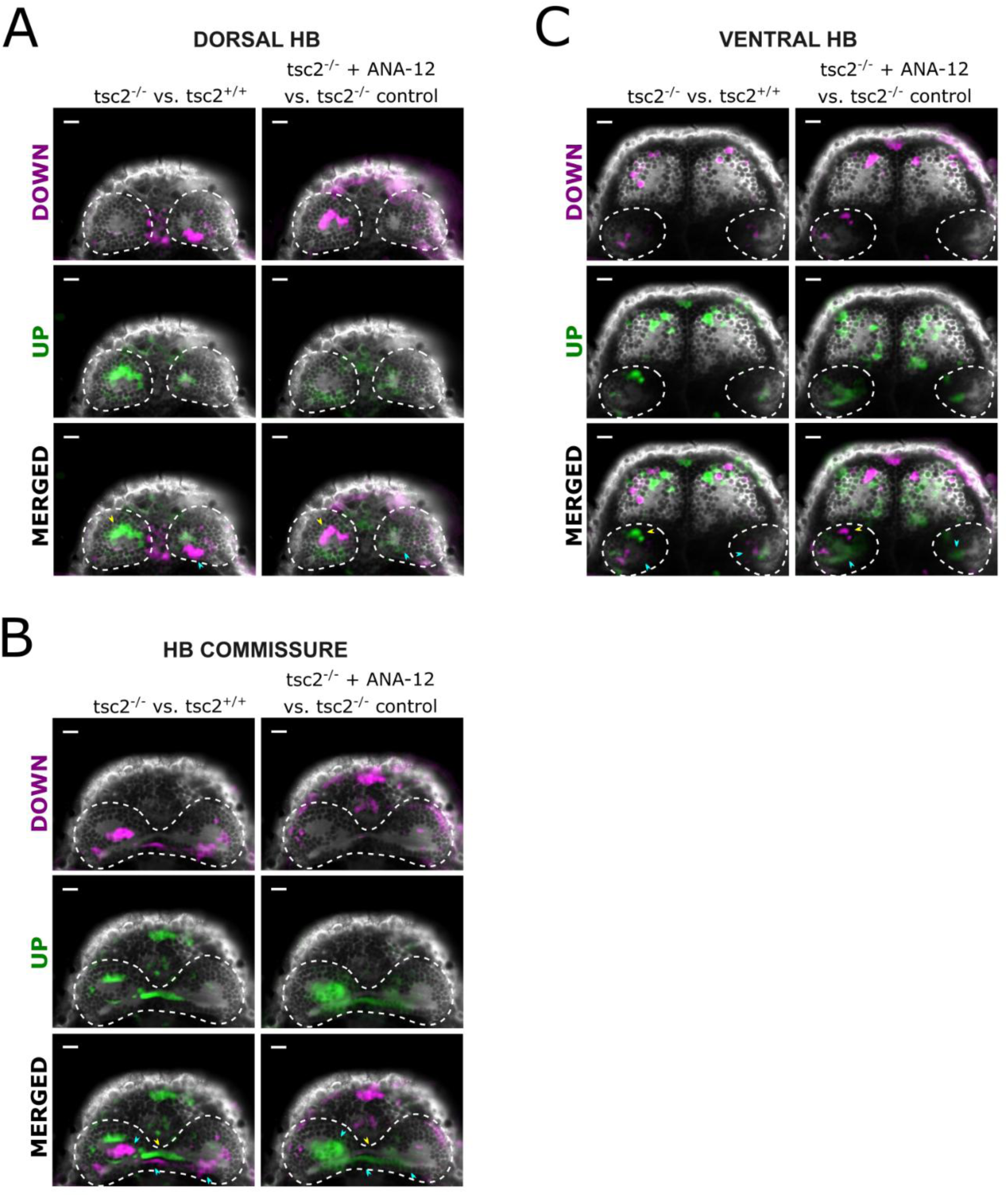
Single z-slices representative of the dorsal habenula (A), area around the habenula commissure (B), and ventral habenula (C). Due to the asymmetry of the habenulae (left Hb being larger than right Hb) the pericommissural area shown here belongs to the dorsal left Hb and ventral right Hb. The images were selected from maps of neuronal activity as measured by the pERK/ERK ratio. Areas marked in magenta represent regions of lower activity in *tsc2^vu242/vu242^* fish vs. wild-type siblings (left), and *tsc2^vu242/vu242^* fish treated with ANA-12 vs. untreated *tsc2^vu242/vu242^* (right), while areas marked in green represent regions of increased activity respectively. The reference brain image is shown in black and white. The yellow and blue arrows point to areas that were upregulated in the mutant but downregulated after ANA-12, or vice versa. Scale bar = 20 µm.

The mammalian amygdaloid complex, located in the forebrain, plays a key role in sensory and behavioral regulation, emotion, and cognition (Pabba, 2013). Most importantly for this study, the amygdala is also crucial for proper responses to fear stimuli (LeDoux, 2000). While its functionality is evolutionarily conserved, a notable difference between zebrafish and mammals is that in the former, the telencephalon forms by eversion, rather than evagination. As a result, the regions of the pallium, while sharing functional homology, are oriented differently than in mammals, making the identification of corresponding structures challenging (Bally-Cuif & Vernier, 2010). While the amygdala has been identified in adult zebrafish (Lal et al., 2018; B. A. Porter & Mueller, 2020), there is still a knowledge gap with regards to amygdala development at early larval stages. We have used the pERK/ERK stainings to assess neuronal activity in the telencephalon of *tsc2^vu242^* fish at 5 dpf, and have stained our samples against a number of markers known to be expressed in various pallial and subpallial territories, in an attempt to identify the amygdaloid nuclei in the developing zebrafish, and correlate the neuronal activity data with functional topology of the brain. Those were the GABAergic neuron marker GAD65/67, the glutamatergic marker VGlut1, neuropeptide Y (NPY), and the calcium-binding proteins calretinin and parvalbumin.

In the most dorsal layer of the *tsc2^vu242/vu242^* zebrafish pallium, we have found both hypo- and hyperactivated cells. Following treatment with ANA-12, most notably, there was an increase of activity along the midline in *tsc2^vu242/vu242^* fish (Fig.4 A). This region was also distinguished by clusters of calretinin-positive cells, located along the medial and posterior edges of the white matter compartments (Fig.4 B-C), although no statistically significant differences in the number of calretinin-positive cells were seen between neither genotypes nor treatment groups (Fig.4 D). Following treatment with ANA-12, sporadic parvalbumin-positive cells were seen in the lateral areas. GAD65/67 and VGlut1 were expressed throughout this layer, with a particularly strong enrichment in the white matter compartments (Fig.4 B).

**Figure 4.**
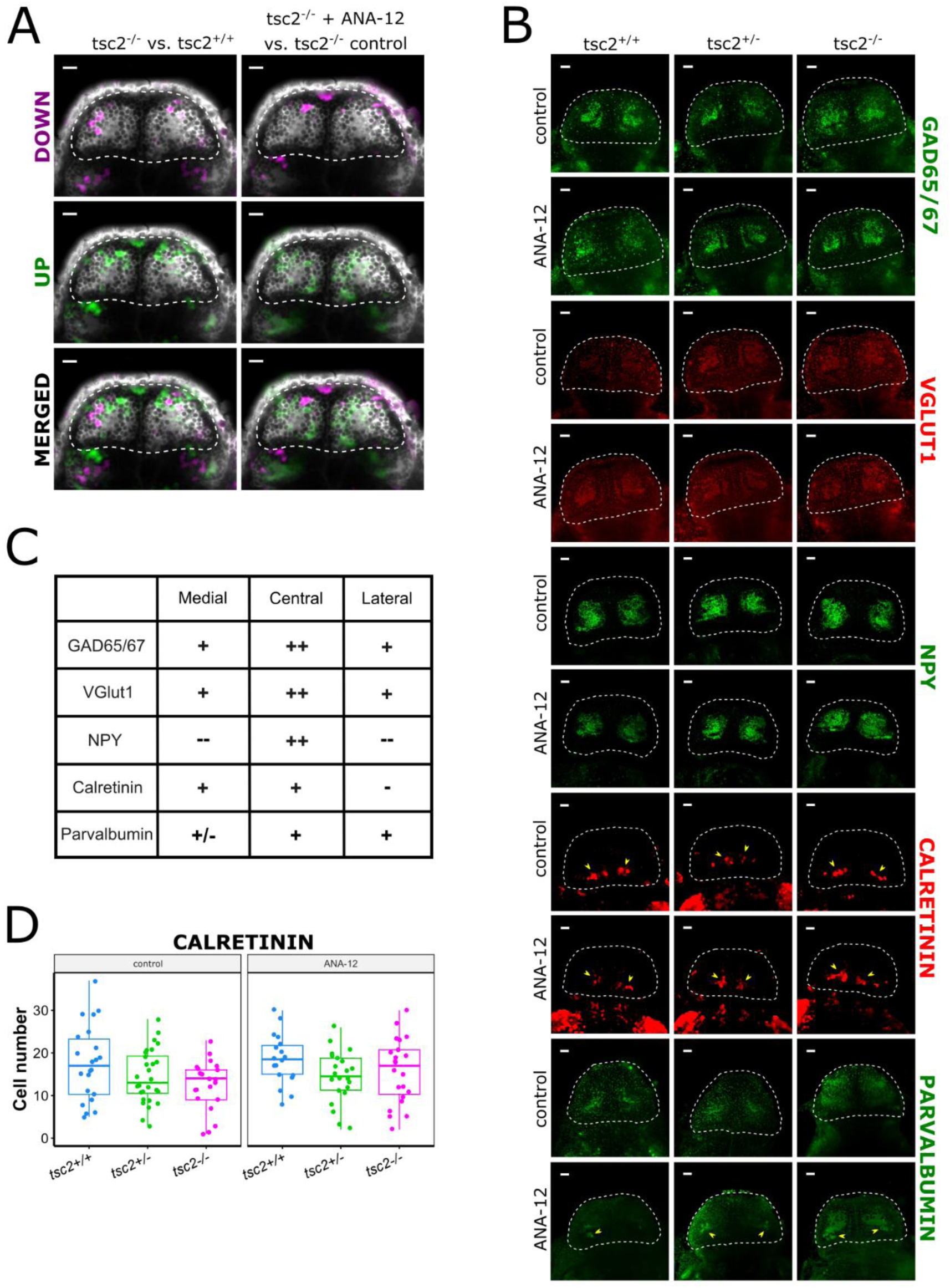
Representative data from the dorsal pallium. (A) Single z-slices from maps of neuronal activity as measured by the pERK/ERK ratio, as described previously. Scale bar = 20 µm. (B) Single z-slices from brains stained with antibodies against GAD65/67, VGlut1, NPY, calretinin and parvalbumin, selected from the corresponding region of the pallium. Yellow arrows point to distinct calretinin- and parvalbumin-positive cells. Scale bar = 20 µm. (C) Expression of markers in the medial, central and lateral areas of this layer, as found in wild-type fish. (D) Number of calretinin-positive cells in the dorsal pallium across all genotypes in untreated vs. ANA-12-treated groups. Each dot represents one fish.

Below this region, moving in the dorsal-ventral axis, multiple hypo- and hyperactivated cells could be seen around the white matter compartments; some of them, though not all, responded to ANA-12 treatment (Fig.5 A). The strong expression of VGlut1/2 together with positive immunoreactivity for GAD65/67 and parvalbumin (Fig.5 A-C) would suggest that this region belongs to the intermediate zone of the subpallium, likely the posterior division of the medial amygdala (MeAp). However, in accordance with data published by Porter and Mueller (B. A. Porter & Mueller, 2020), MeA is distinguished by a high number of calretinin-positive cells, which, in our case, were present only sporadically in this layer (Fig.5 B-C).

**Figure 5.**
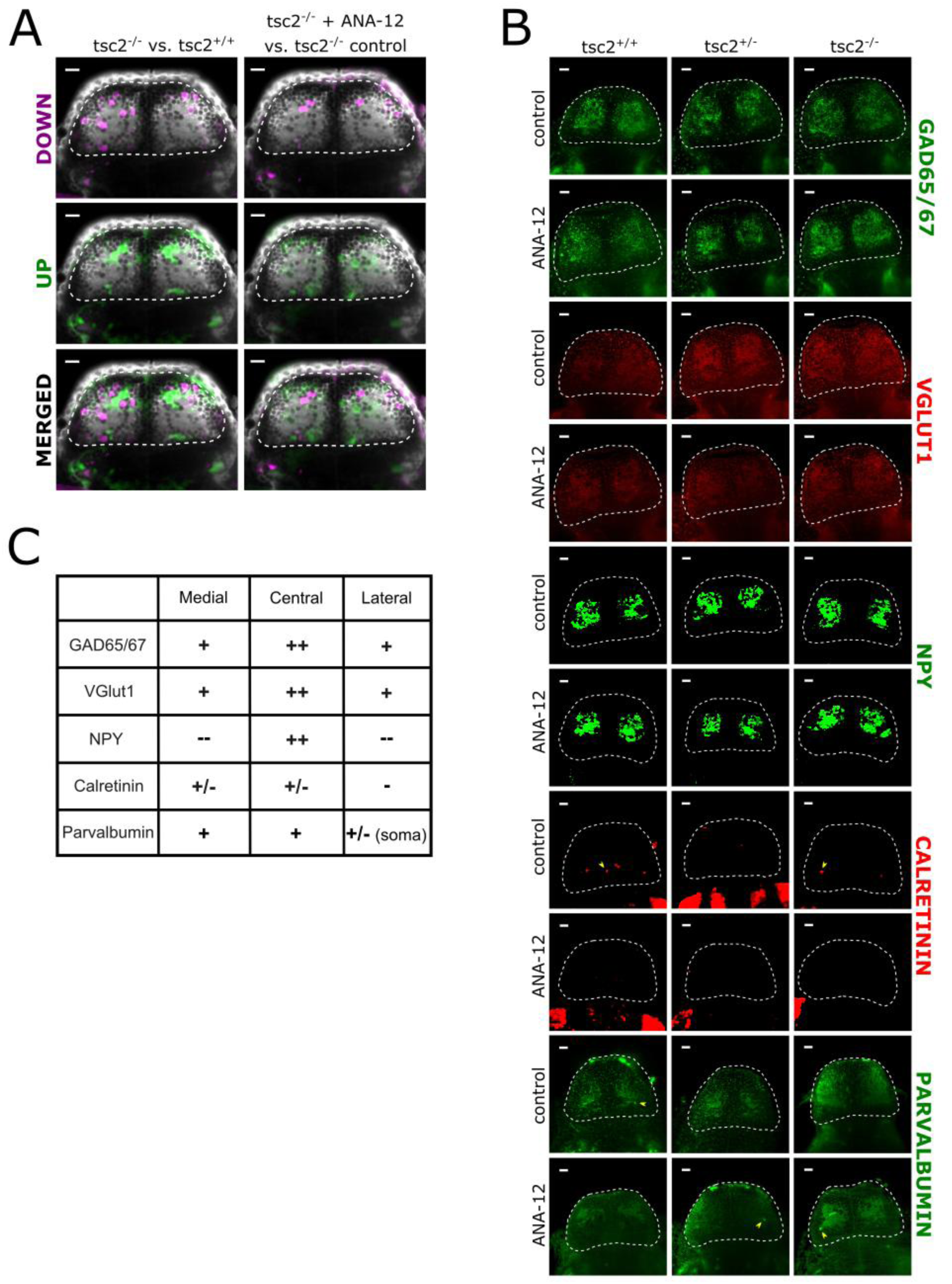
Representative data from the next pallial layer. (A) Single z-slices from maps of neuronal activity as measured by the pERK/ERK ratio, as described previously. Scale bar = 20 µm. (B) Single z-slices from brains stained with antibodies against GAD65/67, VGlut1, NPY, calretinin and parvalbumin, selected from the corresponding region of the pallium. Yellow arrows point to distinct calretinin- and parvalbumin-positive cells. Scale bar = 20 µm. (C) Expression of markers in the medial, central and lateral areas of this layer, as found in wild-type fish.

The next layer in the dorso-ventral axis showed a high increase in neuronal activity in *tsc2^vu242/vu242^* fish compared to wildtype *tsc2^+/+^* siblings (Fig.6 A). Along the brain midline, the neuronal activity was increased even further after ANA-12 treatment, although single hypoactive cells were also seen (Fig.6 A). This area coincided with high numbers of calretinin-positive cells, which was reduced in the *tsc2^vu242/vu242^* mutant, both in the control and treated groups (Fig.6 B-D). In ANA-12-treated fish, there were also downregulated clusters positioned laterally, which overlapped with the localization of parvalbumin-positive cells, and positive immunoreactivity for GAD65/67 (Fig.6 A-C). However, in both cases, it could not be determined for certain if those cells correspond to each other due to a lack of a triple pERK+ERK+parvalbumin or pERK+ERK+calretinin staining. The localization of this region in the dorsal-ventral axis, and the expression of tested markers (GAD65/67+, VGlut1-, calretinin+, parvalbumin+) suggest that this layer might correspond to the anterior division of bed nucleus stria terminalis (BNSTa; labelled as BSTa in Porter & Mueller, 2020), however, this could not be determined for certain. Ultimately, at this stage of development, and with the markers used, we were not able to delineate the precise boundaries between the nuclei of the medial and central amygdala, as well as the BNST.

**Figure 6.**
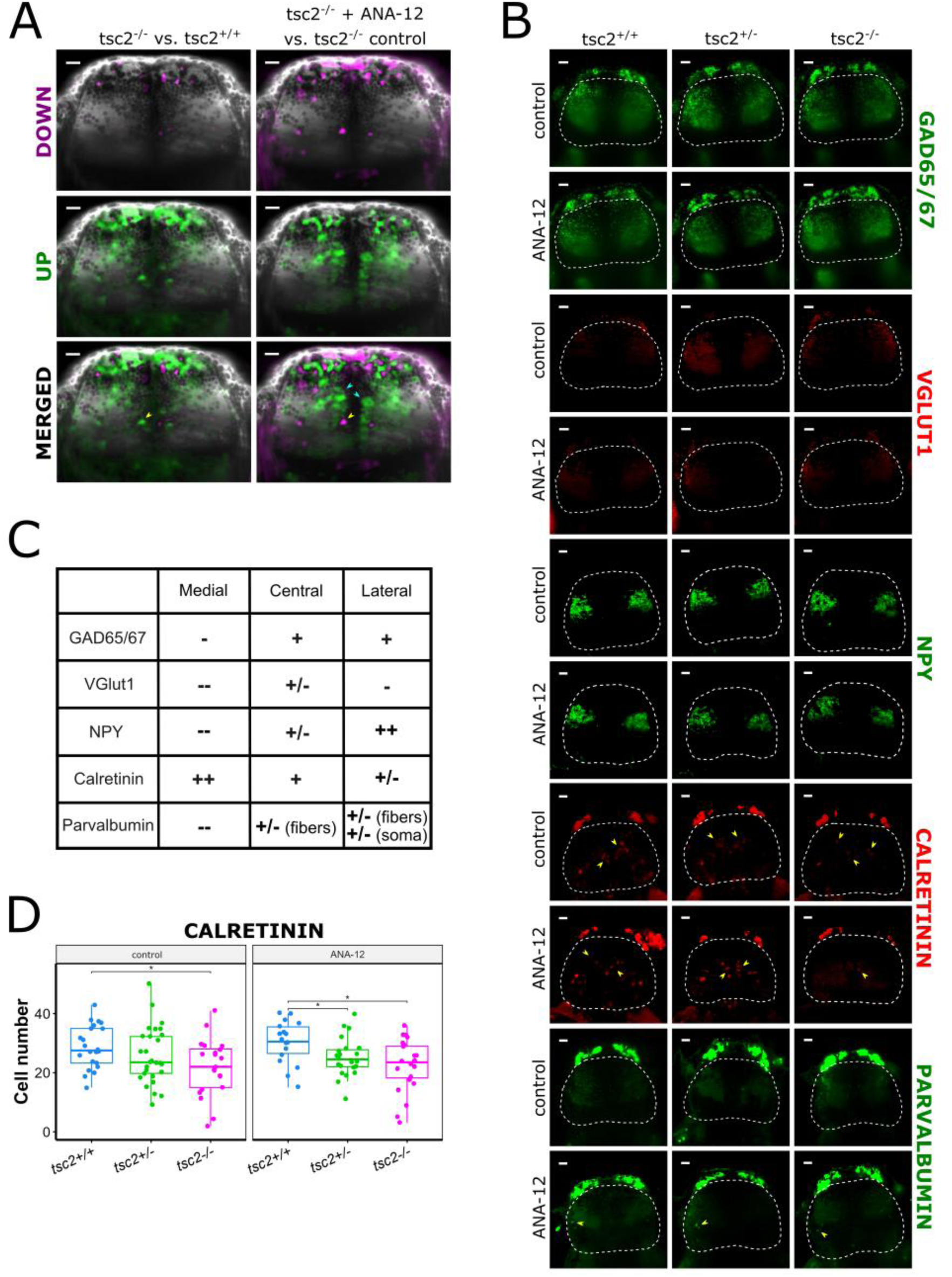
Representative data from the most dorsal layer of the subpallium. (A) Single z-slices from maps of neuronal activity as measured by the pERK/ERK ratio, as described previously. The yellow arrows point to one of the cells that were upregulated in the mutant but downregulated after ANA-12. The blue arrows point to the main area along the midline that was upregulated after ANA-12. Scale bar = 20 µm. (B) Single z-slices from brains stained with antibodies against GAD65/67, VGlut1, NPY, calretinin and parvalbumin, selected from the corresponding region of the pallium. Yellow arrows point to distinct calretinin- and parvalbumin-positive cells. Scale bar = 20 µm. (C) Expression of markers in the medial, central and lateral areas of this layer, as found in wild-type fish. (D) Number of calretinin-positive cells in the subpallium across all genotypes in untreated vs. ANA-12-treated groups [*p* = 0.036 for control *tsc2^+/+^* vs. control *tsc2^vu242/vu242^*, *p* = 0.044 for ANA-12 *tsc2^+/+^* vs. ANA-12 *tsc2^vu242/+^*, *p* = 0.011 for ANA-12 *tsc2^+/+^* vs. ANA-12 *tsc2^vu242/vu242^*]. Each dot represents one fish.

We have also found a population of hyperactive cells in the deeper subpallial layer of *tsc2^vu242/vu242^* telencephalon, which were downregulated by ANA-12 treatment (Fig.7 A). This layer showed negative immunoreactivity for Vglut1. The expression of GAD65/67 and parvalbumin was constrained to the olfactory pallium, where we have also observed a strong calretinin signal (Fig.7 B, D). Single calretinin-positive cells were also found in both the medial and lateral areas of this layer, however, there was no statistically significant difference in their number between neither genotypes nor treatment groups (Fig.7 B-C). Based on its topological position, we have labelled this region as the striatopallidum.

**Figure 7.**
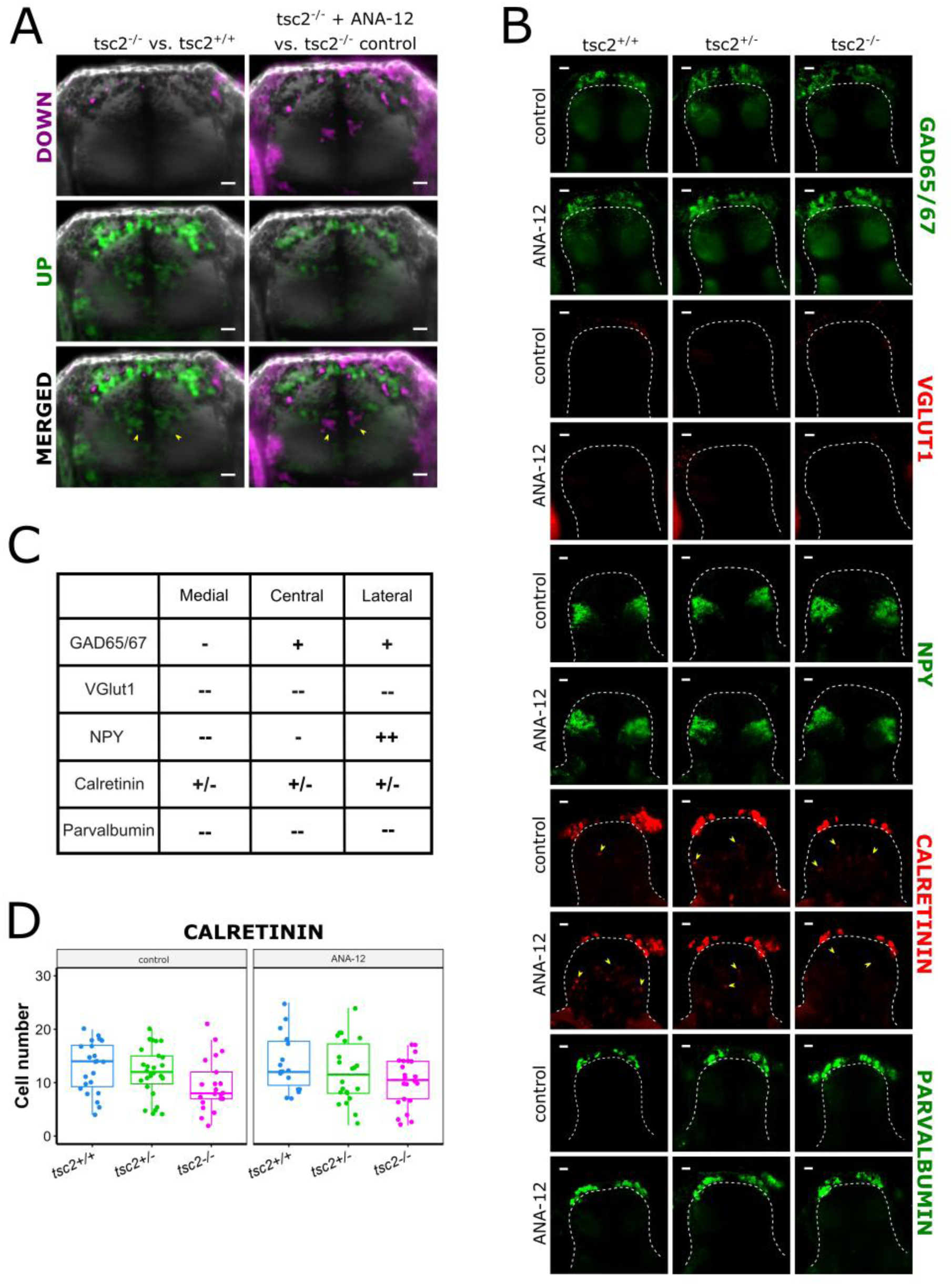
Representative data from the next subpallial layer. (A) Single z-slices from maps of neuronal activity as measured by the pERK/ERK ratio, as described previously. The yellow arrows point to clusters of cells along the midline that were upregulated in the mutant but downregulated after ANA-12. Scale bar = 20 µm. (B) Single z-slices from brains stained with antibodies against GAD65/67, VGlut1, NPY, calretinin and parvalbumin, selected from the corresponding region of the pallium. Yellow arrows point to distinct calretinin-positive cells. Scale bar = 20 µm. (C) Expression of markers in the medial, central and lateral areas of this layer, as found in wild-type fish. (D) Number of calretinin-positive cells in the middle subpallial layer across all genotypes in untreated vs. ANA-12-treated groups. Each dot represents one fish.

The most ventral layer of the *tsc2^vu242/vu242^* fish telencephalon showed strong neuronal hyperactivity in the anterior commissure and the precommissural area (Fig.8 A). By comparison, in *tsc2^vu242/vu242^* fish treated with ANA-12, the centrolaterally positioned regions showed decreased activity, while there was an increase of neuronal activity along the midline (Fig.8 A). We have also found a distinctive V-shaped cluster of parvalbumin-positive cells (Fig.8 B-C). Based on its topological location and immunoreactivity to brain markers, we identified this region as the septum. In the control group, the integrated density of the parvalbumin signal in this area tended towards higher in the *tsc2^vu242/vu242^* mutant, although the difference was not statistically significant. Conversely, in the group treated with ANA-12, the median signal density was higher in the wild-type fish, and significantly lowered in the *tsc2^vu242/vu242^* mutant, compared to both treated wild-type and untreated mutant (Fig.8 C).

**Figure 8.**
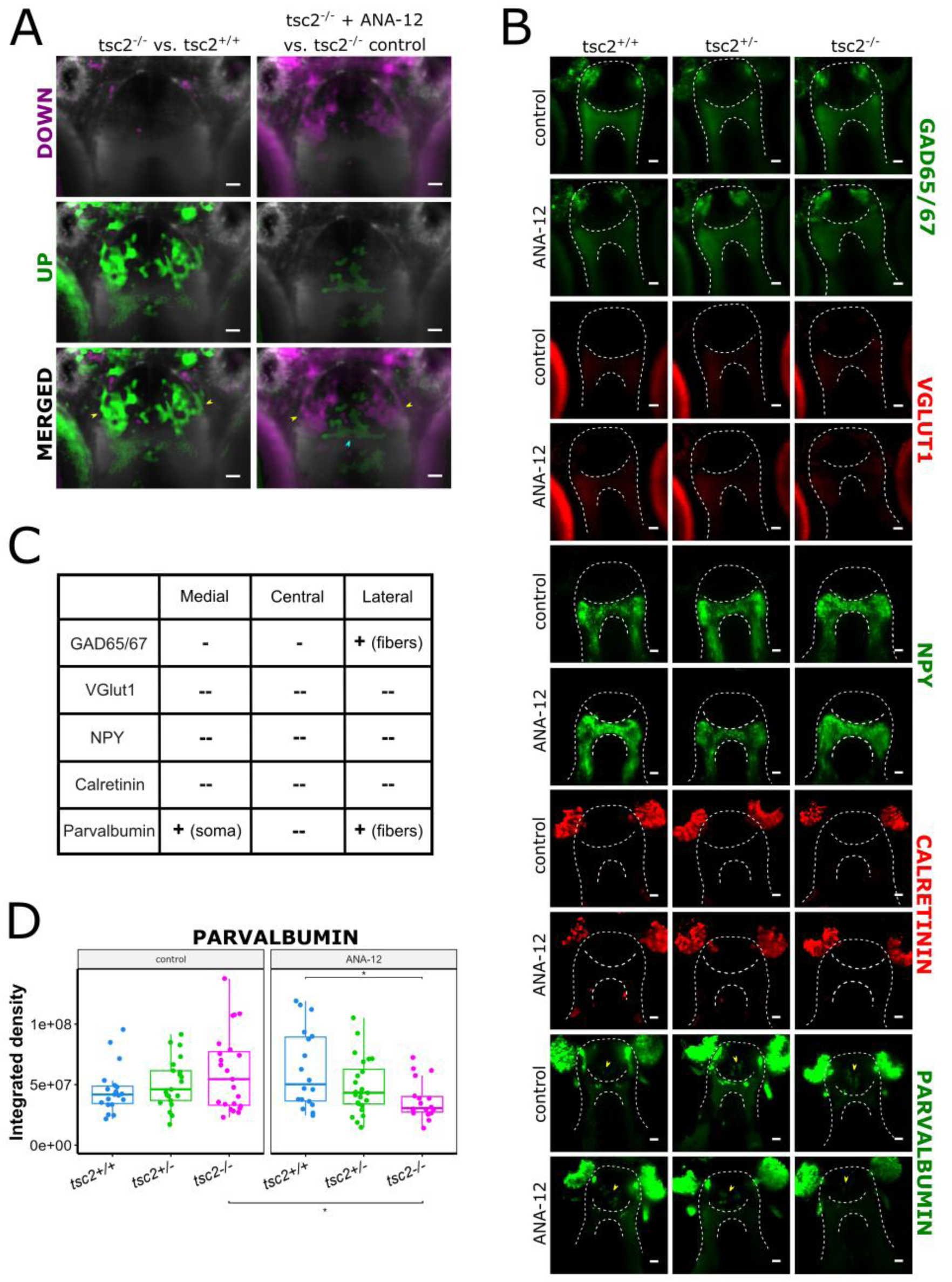
Representative data from the most ventral layer of the subpallium. (A) Single z-slices from maps of neuronal activity as measured by the pERK/ERK ratio, as described previously. The yellow arrows point to laterally positioned areas that that were upregulated in the mutant but downregulated after ANA-12. The blue arrow points to the cluster of cells along the midline that was upregulated after ANA-12. Scale bar = 20 µm. (B) Single z-slices from brains stained with antibodies against GAD65/67, VGlut1, NPY, calretinin and parvalbumin, selected from the corresponding region of the pallium. Yellow arrows point to a distinct cluster of parvalbumin-positive cells. Scale bar = 20 µm. (C) Expression of markers in the medial, central and lateral areas of this layer, as found in wild-type fish. (D) The integrated density count, representing the number and intensity of fluorescent signal, of parvalbumin-positive cells in the ventral subpallium across all genotypes in untreated vs. ANA-12-treated groups [*p* = 0.0129 for control *tsc2^vu242/vu242^* vs. ANA-12 *tsc2^vu242/vu242^*, *p* = 0.032 for ANA-12 *tsc2^+/+^* vs. ANA-12 *tsc2^vu242/+^*]. Each dot represents one fish.

## Discussion

Following our previous findings concerning anxiety-like behavior in *tsc2^vu242/vu242^* fish, and the anxiolytic potential of the TrkB inhibitor ANA-12, our results support the finding that pretreatment with ANA-12 rescues anxiety-like symptoms in *tsc2^vu242/vu242^* larvae. We have confirmed that ANA-12 lowers the hyperactivation of TrkB seen in the *tsc2^vu242/vu242^* mutant, and acts independently of rapamycin-dependent activity of mTorC1. At the same time, we have observed that a 24 h treatment with ANA-12 had the same effect on anxiety-like behavior as treatment with rapamycin from 2 dpf. This suggests that dysregulation of mTorC1 signaling in early development contributes to the etiology of anxiety.

The dorsal habenula is known to regulate fear responses, as experiments with Hb-lesioned fish showed increased freezing behavior in response to conditioned aversive stimuli (Agetsuma et al., 2010; Lee et al., 2010) and increased anxiety in a novel environment (Mathuru & Jesuthasan, 2013) in the lesioned individuals. It was also found through rodent studies that the inhibition of both glutamatergic and GABAergic signaling within the lateral Hb, which corresponds to the ventral Hb in zebrafish, induced anxiety, as measured by exploration in the open field test (Lecourtier et al., 2023). Studies have also demonstrated the role of zebrafish ventral Hb in processing aversive stimuli and learning from negative outcomes (Amo et al., 2014; Chou et al., 2016). Crucially, interrupting the function of ventral Hb affects active avoidance learning, but not panic behavior, which might contribute to exaggerated fear responses as seen in anxiety disorders (Amo et al., 2014). We have found clusters of dysregulated neuronal activity in both dorsal (dHb) and ventral (vHb) habenulae of *tsc2^vu242/vu242^* mutant in comparison to wild-type siblings. The dysregulated neuronal activity that we have observed in the vHb of the *tsc2^vu242/vu242^* mutant might lead to impaired processing of anxiogenic stimuli, and produce a persistent anticipation of negative outcomes, leading to behavioral responses such as reduced exploration in the open field test. Treatment with ANA-12 would then ameliorate the exaggerated anxiety response. Moreover, in our previous work we have noted improper formation of commissural tracts in the *tsc2^vu242/vu242^* mutant, including the anterior (Kedra et al., 2020) and habenular (Doszyn et al., 2024) commissures. The habenula commissure (HC) is formed by tracts projected from the pallium, the eminentia thalami and the posterior tuberculum (Hendricks & Jesuthasan, 2007). Here, we have found that neuronal activity in those tracts was dysregulated in the *tsc2^vu242/vu242^* mutant, potentially contributing to its stunted development, and improper processing of stress-related information by the habenulae. However, further experiments would be necessary to confirm the link between neuronal activity in the HC, habenula dysconnectivity and anxiety-like behavior.

Based on the topological models proposed by Porter and Mueller (B. A. Porter & Mueller, 2020), we attempted to identify components of the amygdaloid complex in the developing zebrafish telencephalon at 5 dpf. We were able to delineate five telencephalic zones, based on their location in the dorsal-ventral axis, as well as the expression patterns of GAD65/67, VGlut1/2, NPY, calretinin and parvalbumin (Fig. 9 A-C). We have putatively labelled them as dorsal telencephalon (D), posterior division of the medial amygdala (MeAp), anterior division of bed nucleus stria terminalis (BNSTa), striatopallidum (Str/PA), and septum (Se).

**Figure 9.**
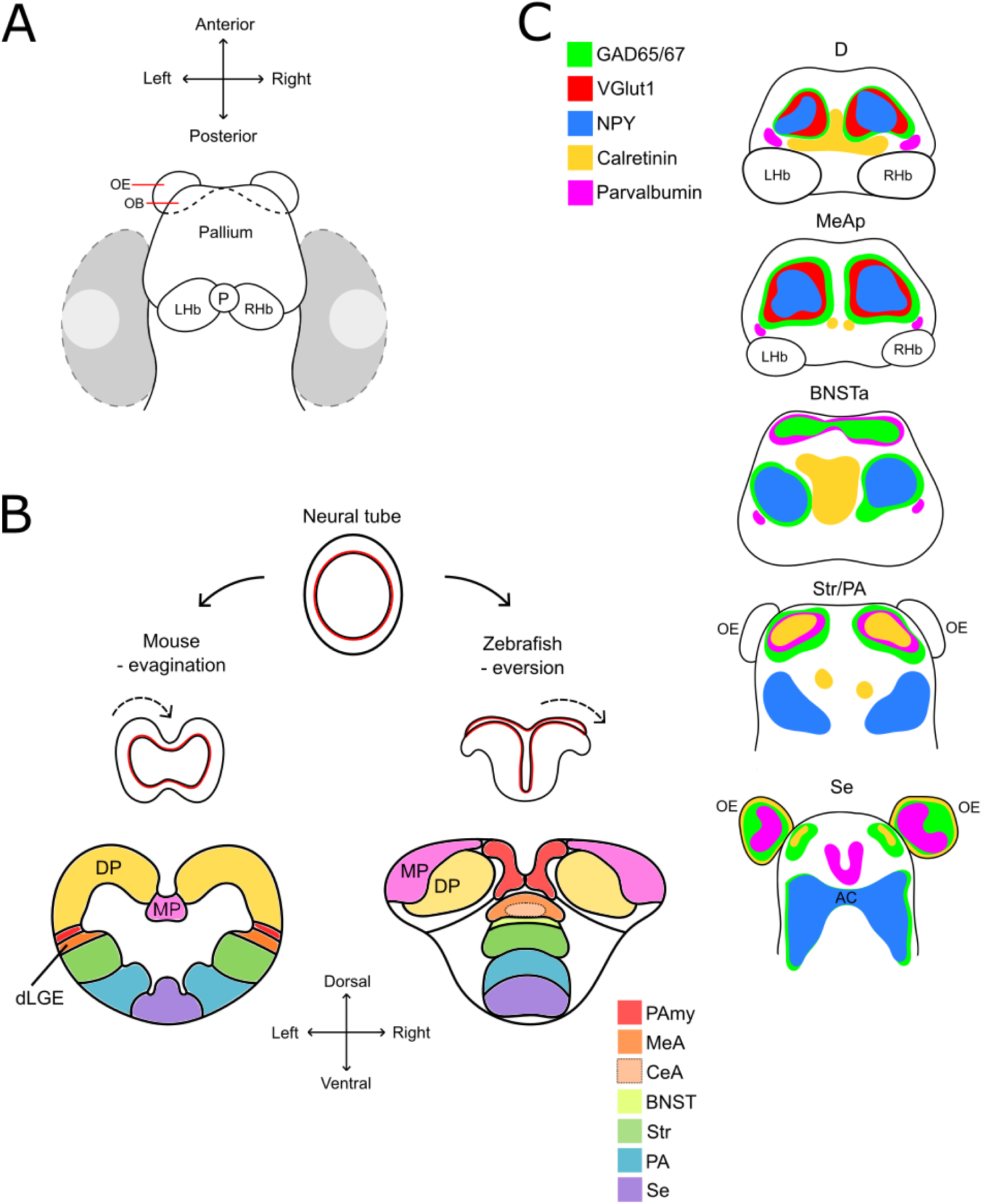
Graphical representations of the developing zebrafish brain. (A) Dorsal view of the forebrain at 5 dpf, showing the locations of the pallium, the olfactory bulbs (OB), olfactory epithelium (OE), pineal complex (P), and left and right habenulae (LHb, RHb), as well as the eyes, marked in grey. This orientation corresponds to the immunofluorescence images shown in the results section. (B) A simplified diagram showing the processes of evagination and eversion, comparing the development of mouse and zebrafish brain, and the location of homologous structures. As seen here, eversion results in the components of the amygdaloid complex (pallial amygdala (PAmy), central and medial amygdala (CeA/MeA), bed nucleus stria terminalis (BNST), striatum (Str), pallidum (PA), and septum (Se)) being located along the midline, rather than laterally. (C) Schematic representations of the localization of GAD65/67, VGlut1, NPY, calretinin and parvalbumin at the level of dorsal telencephalon (D), posterior division of MeA (MeAp), anterior division of BNST (BNSTa), striatopallidum (Str/PA) and Se, based on our immunofluorescence data.

The basolateral amygdala (BLA) plays a crucial role in fear conditioning (LeDoux, 2000). Glutamatergic neurons found in the basolateral amygdala (BLA) are key to fear learning following aversive stimuli, and their inhibition impairs this process (Sengupta et al., 2018). The majority of GABAergic interneurons in the BLA express calcium-binding proteins parvalbumin or calretinin, though cells expressing those markers form separate populations (Sah et al., 2003). Studies on goldfish, and later also zebrafish, pointed to the dorsomedial telencephalon as its teleostan equivalent (Aoki et al., 2013; Portavella et al., 2002), and it is currently believed that the zebrafish central and pallial amygdala territories correspond to the mammalian BLA (B. A. Porter & Mueller, 2020). It is possible that in our horizontal sections of the brain, the calretinin-positive cells in the first dorsal layer belong to the pallial amygdala, however, we have not been able to confirm this; we have also not subjected our larvae to repeated aversive stimuli to determine whether they also experience a dysregulation in fear learning.

While we were not able to precisely define the boundaries of pallial, central and medial amygdala at this stage of development, we have identified the localization Str/PA and Se. We based our identification on their location in the dorsal-ventral axis, by comparing our results to previously published models of the zebrafish telencephalon (Gerlach & Wullimann, 2021; B. A. Porter & Mueller, 2020; Tanimoto et al., 2024). These ventral regions of the subpallium were also where we have noted the most prominent differences in neuronal activity between *tsc2^vu242/vu242^* larvae and wild-type siblings, that were also clearly reversed by ANA-12 treatment.

In humans, various parts of the striatum are involved in decision making and motivation (J. N. Porter et al., 2015). Rodent studies have also shown that dopaminergic neurons in the nucleus accumbens (NAc) and the posterior tail of the striatum (TS) take part in threat learning (Duvarci, 2024). Notably, dopaminergic projections into the TS drive avoidant behaviors in response to novel stimuli, but do not encode outcome values (Menegas et al., 2018). Therefore, it is possible that the hyperactivity of striatal neurons in *tsc2^vu242/vu242^* fish is similarly linked to exaggerated threat avoidance and “anticipatory” anxiety that is not ameliorated by a lack of actual negative outcomes.

In mammals, the lateral septum (LS) is comprised of both anxiolytic and anxiogenic neuronal populations (Chen et al., 2021; Rizzi-Wise & Wang, 2021). The heterogeneity of LS function could also be related to its role in assessing changes in valence. It has been hypothesized that the LS might integrate motivation, mood and movement signals. Therefore, animals with dysregulated LS would respond incorrectly to various environmental cues, and could react with exaggerated motor responses (Wirtshafter & Wilson, 2021). The medial septum (MS) is also known to be involved in regulation of anxiety (Chang et al., 2025), as well as processing fear memory (Calandreau et al., 2007). Specifically, inhibiting the function of MS resulted in impaired extinction of fear memory (Knox & Keller, 2016; Tomaszewski et al., 2024; Tronson et al., 2009). While we have not delineated sub-regions of the septum, the lateral regions which were hyperactivated in *tsc2^vu242/vu242^* fish and downregulated by ANA-12 could correspond to the parts of mammalian LS that promote anxiety-like behavior. At the same time, we have observed an upregulation of the medial septum in fish treated with ANA-12. However, additional behavioral testing would be necessary to determine whether ANA-12 promotes boldness and exploratory behavior by increasing fear memory extinction.

In summary, we have found that multiple brain regions involved in anticipation of negative outcomes and processing anxiogenic stimuli are dysregulated in the *tsc2^vu242/vu242^* mutant. Treatment with ANA-12 rescued this impairment in multiple areas, particularly the habenula and ventral subpallium. We were also able to identify the latter as corresponding to the mammalian striatopallidum and septum. A further investigation into the neurotransmitters released by the affected neurons and their connectivity could resolve the precise mechanism of TrkB-regulated anxiety-like behavior in the *tsc2^vu242/vu242^* zebrafish model.

### Limitations

In the larval zebrafish, such as were used in this study, the brain is smaller, and the processes of eversion and outgrowth are still in progress, which makes distinguishing between various brain regions more challenging than in adults. Additionally, some markers might not yet be expressed in early development, or be expressed differently because cells are not yet fully specified. Nevertheless, further studies including more brain markers should help in distinguishing regions that we were not able to identify. Other than by spatial inference, we were also not able to directly correlate neuronal activity with the expression of marker proteins, as the limited availability of zebrafish-specific antibodies did not allow for triple stainings with pERK/ERK. Therefore, for neurons with observed differences in activity, we were not able to fully classify them by neurotransmitter or molecular markers.

## Resources availability

### Lead contact

Further information and requests for resources and reagents should be directed to and will be fulfilled by the Lead Contact, Justyna Zmorzynska.

### Materials availability

This study did not generate new unique reagents. The fish mutant and transgenic lines are protected under material transfer agreement with the institutions that generated the lines. Upon appropriate agreement with these institutions, they can be requested from the lead contact.

### Data and code availability

Data: All data is included in the manuscript. Microscopy data reported in this paper will be shared by the lead contact upon reasonable request.

Code: This paper does not report original code.

Additional information: Any additional information required to reanalyze the data reported in this paper is available from the lead contact upon request.

## Acknowledgements

We thank Kevin Ess (Vanderbilt University) for the *tsc2^vu242/+^* zebrafish line, the IIMCB Zebrafish Core Facility for assistance with the adult fish, and the IIMCB Microscopy Facility for sharing the Lightsheet Z.1. This work was supported by an OPUS grant no. 2020/37/B/NZ3/02345 (J.Z.) from National Science Centre, Poland.

## Author contributions

O.D.: investigation, formal analysis, visualization, writing – original draft, writing – review & editing; J.Z.: conceptualization, investigation, formal analysis, writing – review & editing, supervision, project administration, funding acquisition.

